# Stroke survivors show task-dependent modulation of motor variability during bimanual coordination

**DOI:** 10.1101/292193

**Authors:** Rajiv Ranganathan, Rani Gebara, Michael Andary, Jim Sylvain

## Abstract

Stroke often results in hemiparesis, leaving one side of the body ‘affected’ relative to the other side. Prior research has shown that the affected arm has higher variability – however, the extent to which this variability can be modulated is unclear. Here we used a shared bimanual task to examine the degree to which participants could modulate the variability in the affected arm after stroke. Participants with chronic stroke (n = 11), and age-matched controls (n = 11) performed unimanual and bimanual reaching movements to move a cursor on a screen to different targets. In the unimanual condition, the cursor was controlled only by the movement of a single arm whereas in the bimanual condition, the cursor position was “shared” between the two arms by using a weighted average of the two hand positions. Unknown to the participants, we altered the weightings of the affected and unaffected arms to cursor motion and examined how the movement variability on each arm changed depending on its contribution to the task. Results showed that stroke survivors had higher movement variability on the affected arm – however, like age-matched controls, they were able to modulate the variability in both the affected and unaffected arms according to the weighting condition. Specifically, as the weighting on a particular arm increased (i.e. it became more important to the task), the movement variability decreased. These results show that stroke survivors are capable of modulating variability depending on the task context, and this feature may potentially be exploited for rehabilitation paradigms

## 1 Introduction

Hemiparesis in the upper extremity after stroke, which leaves one side of the body more ‘affected’ (also referred to as the paretic side) compared to the other ‘unaffected’ (non-paretic) side, can cause significant difficulty with performing activities of daily living (Nakayama et al., 1994). Deficits in the affected arm after stroke have been characterized extensively and include muscle weakness, impaired coordination, and higher motor variability (Dewald et al., 1995; Ada et al., 2003; Cirstea et al., 2003; Krakauer, 2005; Lang et al., 2005). Developing new methods to address these deficits remains a key challenge in stroke rehabilitation.

However, while a predominant number of studies have compared the affected arm and unaffected arm in unimanual tasks (i.e. when only one arm is moving at a time), there have also been several studies which examine the interaction between the arms during bimanual tasks (Cunningham et al., 2002; Rose and Winstein, 2005; Metrot et al., 2013; Gosser and Rice, 2015; Kantak et al., 2016). Bimanual tasks are of interest not only from a functional standpoint (as many activities of daily living involve bimanual movements), but are also interesting from a theoretical standpoint because in addition to quantifying the deficits of the affected arm, they provide insight into whether the coordination between the arms is disrupted after stroke. Understanding the interaction between the two arms to facilitate the recovery of the affected arm has been the focus of bimanual therapy in stroke (Whitall et al., 2000; Luft et al., 2004; Cauraugh et al., 2010; Kantak et al., 2017)

One specific type of bimanual task that is particularly of interest is where the two arms ‘share’ the same task goal (e.g. holding a tray level with both hands). In this case, a feature of the task is motor redundancy (Bernstein, 1967) – i.e., one arm can ‘compensate’ directly for deficits in the other arm (Latash et al., 2002; Diedrichsen, 2007). A specific prediction from the optimal feedback control framework is that in such redundant tasks control is ‘flexible’ – i.e. the nervous system does not always control all variability to a minimum– but rather allows variability as long as it does not influence task performance (Todorov and Jordan, 2002; Diedrichsen and Dowling, 2009). Prior research in redundant isometric force production tasks has shown that stroke survivors had higher asymmetries in force production, and reduced coordination between the two arms (Lodha et al., 2012a, 2012b). However, because these studies only examined a single ‘shared’ condition (where the total force was the sum of the forces exerted by the two arms), the degree to which this variability in the affected arm can be modulated by stroke survivors is still unclear.

In the current study, we examined if stroke survivors are capable of modulating the variability in the affected and unaffected arms depending on the task. We used a virtual reaching task where participants had to control a ‘shared cursor’ that was controlled by the motion of both arms (Diedrichsen, 2007; Mutha and Sainburg, 2009). Specifically, we altered the contribution of each arm to the task to examine if the variability in each arm is affected by the task demands. Our hypothesis was that stroke survivors would show higher variability in the affected arm compared to the non-affected arm, but be able to modulate this variability specific to the task context.

## 2 METHODS

### 2.1 Participants

There were two groups of participants (n = 22 total). The stroke group (n = 11, 8 males, age range – 29 to 67 years) consisted of chronic stroke survivors. Inclusion criteria were (i) the occurrence of a stroke at least 9 months prior to participation, and (ii) ability to make reaching movements with both hands to targets that were placed on a table (ability to grasp was not required). Exclusion criteria were visual or cognitive deficits that interfered with viewing the cursor or understanding task instructions. The control group was a group of individuals who were roughly age-matched (n = 11, 7 males, age range – 32 to 70 years) who had no history of conditions that affected their ability to move their arms. Participants provided informed consent and were compensated for their time, and all experimental procedures were approved by the Human Research Protection Program at Michigan State University.

### 2.2 Experimental Setup

A 4-camera motion capture system (Motion Analysis Corporation) was used to capture the 3D positions of various landmarks on the upper body. Ten retroreflective markers were placed on the head, sternum, and bilaterally on the shoulders, elbows, wrists, and the hands (third metacarpal joint) (Fig. 1). The sampling rate of the cameras was set at 120 Hz.

**Figure 1.**
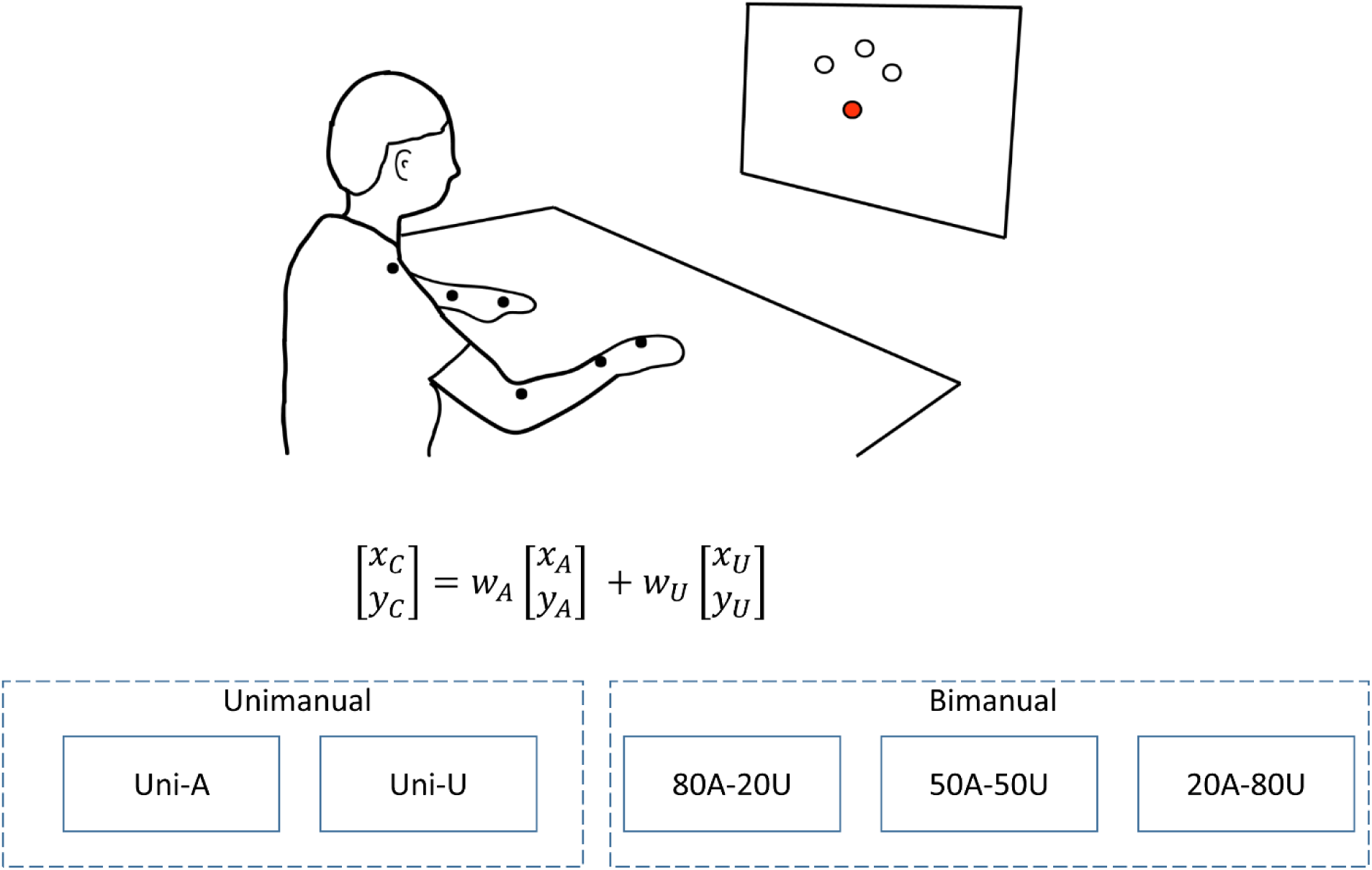
Experimental Setup. Retroreflective markers were attached to the major joints of the upper body and tracked with a motion capture system. The hand positions in real-time were used to display a cursor on the screen, which had to be moved to three targets. There were two types of conditions - in the unimanual conditions, only the affected (Uni-A) or the unaffected hand (Uni-U) controlled the cursor; in the bimanual conditions, the cursor position was a weighted average of the two hand positions. The weighting of the two hands was varied from greater weighting on the affected hand (80A-20U), equal weighting on both hands (50A-50U), and greater weighting on the unaffected hand (20A-80U). Participants completed all trials of one condition before moving to the other conditions.

### 2.3 Terminology

We would like to clarify the nomenclature used throughout the rest of the manuscript. In the stroke population, the ‘affected arm’ was the paretic arm, and the ‘unaffected arm’ was the non-paretic arm (regardless of arm dominance). For the age-matched controls, the ‘affected arm’ was the non-dominant arm, and the ‘unaffected arm’ was the dominant arm. Arm dominance was self-reported by participants.

### 2.4 Task

Participants performed a virtual reaching task which required them to move one or both of their arms on a table to move a cursor to set of targets displayed on a computer monitor. There were three targets placed on the screen at a distance of 15 cm from the home position (Fig. 1). Participants were instructed to reach to the target as quickly and as accurately as possible. A scoring system was provided to encourage fast reaching. Each trial was considered complete only when the previous target was successfully reached.

There were two types of reaching conditions – In the unimanual conditions, the cursor position was controlled only by position of the affected hand (Uni-A), or the unaffected hand (Uni-U). This meant that there was no redundancy in unimanual trials.

In the bimanual conditions, the cursor position ***X***_***C***_ (which denotes the 2-D x,y position) was determined as a weighted average of the affected and unaffected hand positions (***X***_***A***_ and ***X***_***U***_ respectively) as follows:

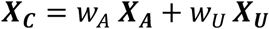

Where w_A_, w_U_ were the weighting factors that determined the contribution from each limb. In the 50A-50U condition, (w_A_,w_U_) = (0.5,0.5), and the cursor was exactly at the midpoint between the two hands with equal contribution from both hands. In the 80A-20U condition, (w_A_,w_U_) = (0.8,0.2), which meant that the cursor was much more heavily weighted on the affected hand compared to the unaffected hand. Finally, in the 20A-80U condition, (w_A_,w_U_) = (0.2,0.8) and the cursor was much more heavily weighted on the unaffected hand compared to the affected hand. By adjusting these weighting factors, we systematically altered the contribution of each hand to the cursor control to examine whether the coordination between the arms was sensitive to these task demands.

### 2.5 Procedure

After providing informed consent, participants performed a brief familiarization phase with both unimanual and bimanual reaches to make sure they understood the task. After this familiarization phase, each participant performed 5 blocks of reaching (Fig. 1). There were 2 unimanual reaching blocks, where participants only reached with either the paretic (non-dominant arm for controls) or the non-paretic arm (dominant arm for controls). There were also 3 bimanual reaching blocks, where the weights were either: 80A-20U, 50A-50U or 20A-80U. In the unimanual conditions, each block consisted of 30 reaches (10 to each of the three targets), whereas in the bimanual conditions each block consisted of 45 reaches (15 to each of the three targets). All trials in a block were performed consecutively. The order of the 5 blocks were randomized with the constraint that participants completed all three bimanual blocks before completing the two unimanual blocks (or vice versa). During the unimanual blocks, participants were instructed to reach only with one arm. During the bimanual reaching blocks, participants were instructed to reach with both arms, but importantly, no information was provided to the participant about the weighting of each arm.

## 3 DATA ANALYSIS

### 3.1 Movement Time

We computed the movement time of each reach as the time between the cursor left the home position, and when it entered the target (and did not leave the target). These instants were determined using a distance threshold of 0.1 cursor units (which corresponded approximately to 7.5 mm).

### 3.2 Spatial variability

The key variables of interest were the spatial variabilities of the cursor, affected arm, and the unaffected arm during the reach. The 2-D position of each reaching movement (which included 2-D position of the cursor, the affected and unaffected hands) were segmented into equispaced intervals of 25% with 0% being the home position, and 100% being the point when the cursor landed in the target. For each block of trials, the spatial variability was computed orthogonal to the direction of the target (Todorov and Jordan, 2002; Ranganathan and Newell, 2010).

### 3.3 Correlation between left and right arms

For the three bimanual conditions, we also computed a correlation between the two arms to examine the degree to which the arms cooperated during the task. Specifically we computed the correlation between the deviations from the mean trajectory in the two hands (in the direction orthogonal to the direction of the target). In this metric, a highly negative correlation would imply that the two arms are cooperating with each other (i.e. when one hand moved to the left, the other hand moved to the right thereby keeping the cursor relatively at the same position) – on the other hand a correlation close to zero would imply that deviations in both arms were independent of each other.

## 4 STATISTICAL ANALYSIS

Movement time was analyzed using a 5 × 2 (Condition × Group) ANOVA, with condition (Uni-A, 80A-20U, 50A-50U, 20A-80U, Uni-U) as the within-subject factor, and group (stroke, control) as the between-subject factor.

Both the spatial variability of each hand and the correlations were analyzed using an n × 5 × 2 (Condition × Phase × Group) mixed model ANOVA with condition and phase as the within-subject factor, and group (stroke/control) as the between-subject factor. The number of conditions (n) in the ANOVA changed depending on the dependent variable – all 5 conditions were included for the analysis of the cursor variability, 4 conditions were included for the affected/unaffected arm variability (because the affected/unaffected arm was not used in one of the unimanual conditions), and 3 conditions were included for the correlation between the two arms (because the correlation could only be computed in the bimanual conditions).

Post-hoc comparisons were performed using method of simple main effects, and the Greenhouse-Geisser correction was applied whenever there was a violation of sphericity. The level of significance was set at α = .05.

## 5 RESULTS

Movement trajectories from a representative participant in all 5 conditions are shown (Fig. 2A).

**Figure 2.**
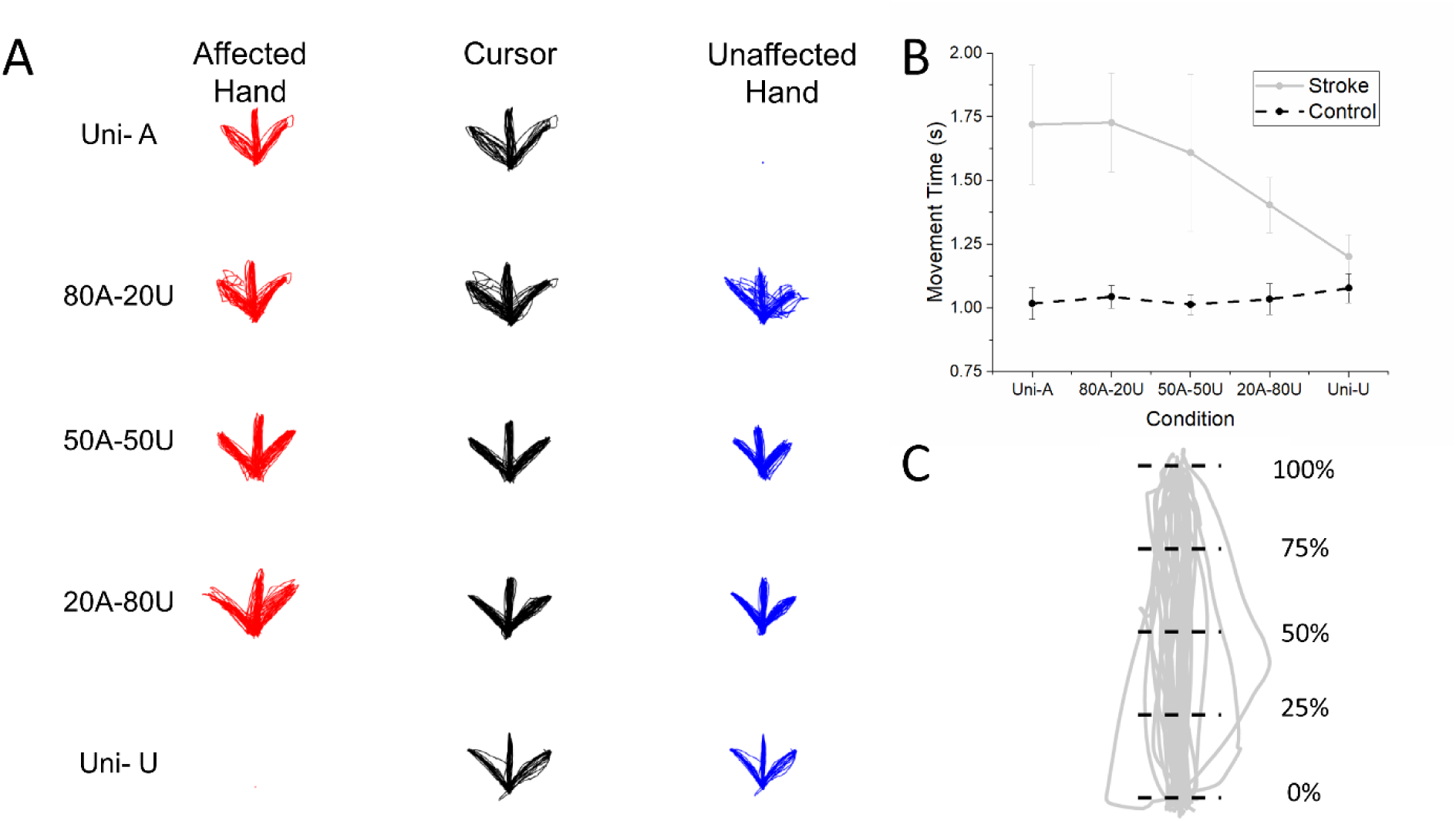
(A) Sample trajectories of the affected hand (red), the unaffected hand (blue), and the cursor (black) from one participant in the stroke group in the 5 conditions. Participants showed modulation of variability depending on the task demands. Specifically, participants increased the motor variability when the contribution of the hand to the cursor position was lower, and decreased the variability of when the contribution of the hand to the cursor position was higher. (B) Movement time for both groups across the different conditions. Error bars represent one SEM, (C) Schematic of spatial variability calculation – trajectories were split into 5 equispaced intervals from the home (0%) to the target (100%), and the standard deviation perpendicular to the direction of motion was measured.

### 5.1 Movement Time

The analysis of movement time indicated that movement time differed between group (Fig 2B). There was a significant main effect of group, F(1,20) = 9.055, p = .007, with stroke survivors showing longer movement times compared to controls. There was also a marginally significant condition × group interaction, F(2.24, 44.73) = 2.835, p = .064, which indicated that stroke survivors had higher movement times for conditions with greater weightings on the affected arm, but controls did not show such modulation. The main effect of condition, F(2.24, 44.73) = 2.026, p = .139 was not significant.

### 5.2 Spatial Variability

A schematic of the spatial variability computation is shown (Fig. 2C).

#### Cursor variability

The analysis of cursor variability indicated that the variability was modulated along the path, and was influenced by the condition (Fig. 3). There was a main effect of phase F(1.22,24.38) = 48.378, p < .001, and a main effect of condition, F(2.24,44.77) = 3.254, p = .043. Post hoc analysis of the phase effect showed an inverted-U shaped trend where the cursor variability was lowest at the start and end of the movement (i.e., 0% and 100% into the path) and highest during the middle of the movement (i.e. at 50% into the path). Post hoc analysis of the condition effect showed that the Uni-A condition had higher variability than all other conditions except the 80A-20U condition. All other main effects and interactions were not significant.

**Figure 3.**
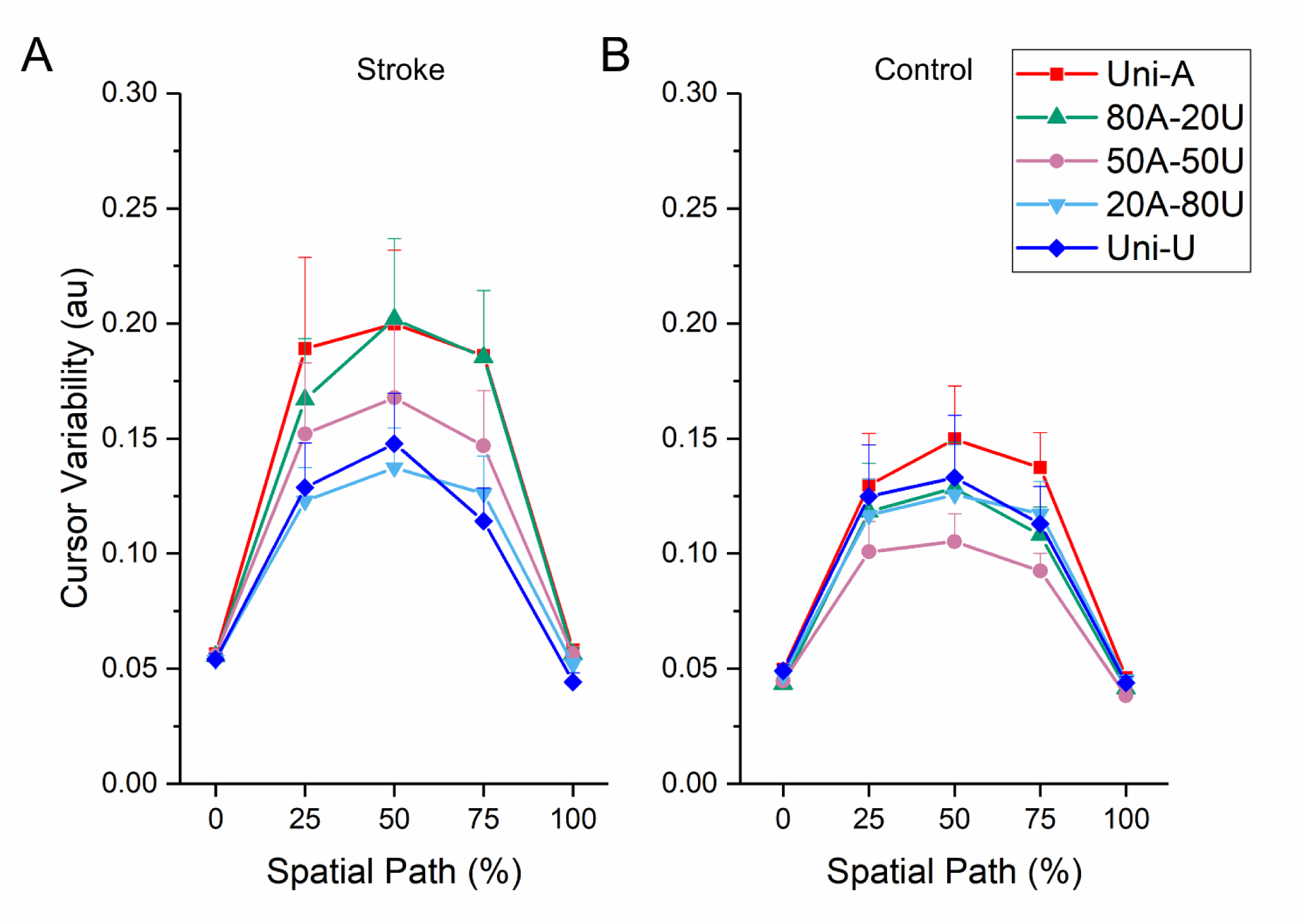
Variability of the cursor along different points along the spatial path in (A) stroke and (B) control groups. The spatial path was split into 5 equally spaced intervals starting from 0% (home position) to 100% (when the cursor reached the target). Both the stroke and control groups showed similar modulation of variability along the path in all conditions, with the highest variability at 50% - i.e. roughly halfway along the distance between the home position and the target.

#### Affected hand variability

The analysis of the variability in the affected hand also showed a significant modulation of variability along the path and was influenced by the condition, but here we found that the modulation of variability depended on the condition. There was a main effect of phase F(1.93, 38.64) = 25.085, p<.001, a main effect of condition, F(1.98,39.67) = 24.678, p<.001, which were mediated by a significant Phase × Condition interaction F(2.96,59.10) = 11.861, p<.001. Post-hoc analysis of the interaction showed that the while conditions with higher weighting on the affected arm (Uni-A and 80A-20U) showed the inverted-U pattern in the variability (that was also seen in the cursor variability), conditions with lower weighting (50A-50U, 20A-80U) did not show this pattern. There was also a significant main effect of group, F(1,20) = 8.692, p = .008, with the stroke group showing higher variability than the control group.

#### Unaffected hand variability

The analysis of the variability in the unaffected hand also showed similar effects to the affected hand. There was a significant modulation of variability along the path and was influenced by the condition, but once again the modulation of variability along the path depended on the condition. There was a main effect of phase, F(1.54, 30.82) = 20.671, p < .001, a main effect of condition, F(2.00,40.03) = 13.735, p <. 001, which was mediated by a significant Phase × condition interaction F(5.16,103.15) = 9.679, p<.001. Post-hoc analysis of the interaction showed that while conditions with higher weighting on the unaffected arm (Uni-U and 20A-80U) showed the inverted-U pattern in the variability (that was also seen in the cursor variability), conditions with lower weighting (50A-50U, 80A-20U) did not show this pattern. The effect of group was marginally significant, F(1,20) = 4.072, p = .057, with the stroke group showing higher variability than the control group.

**Figure 4.**
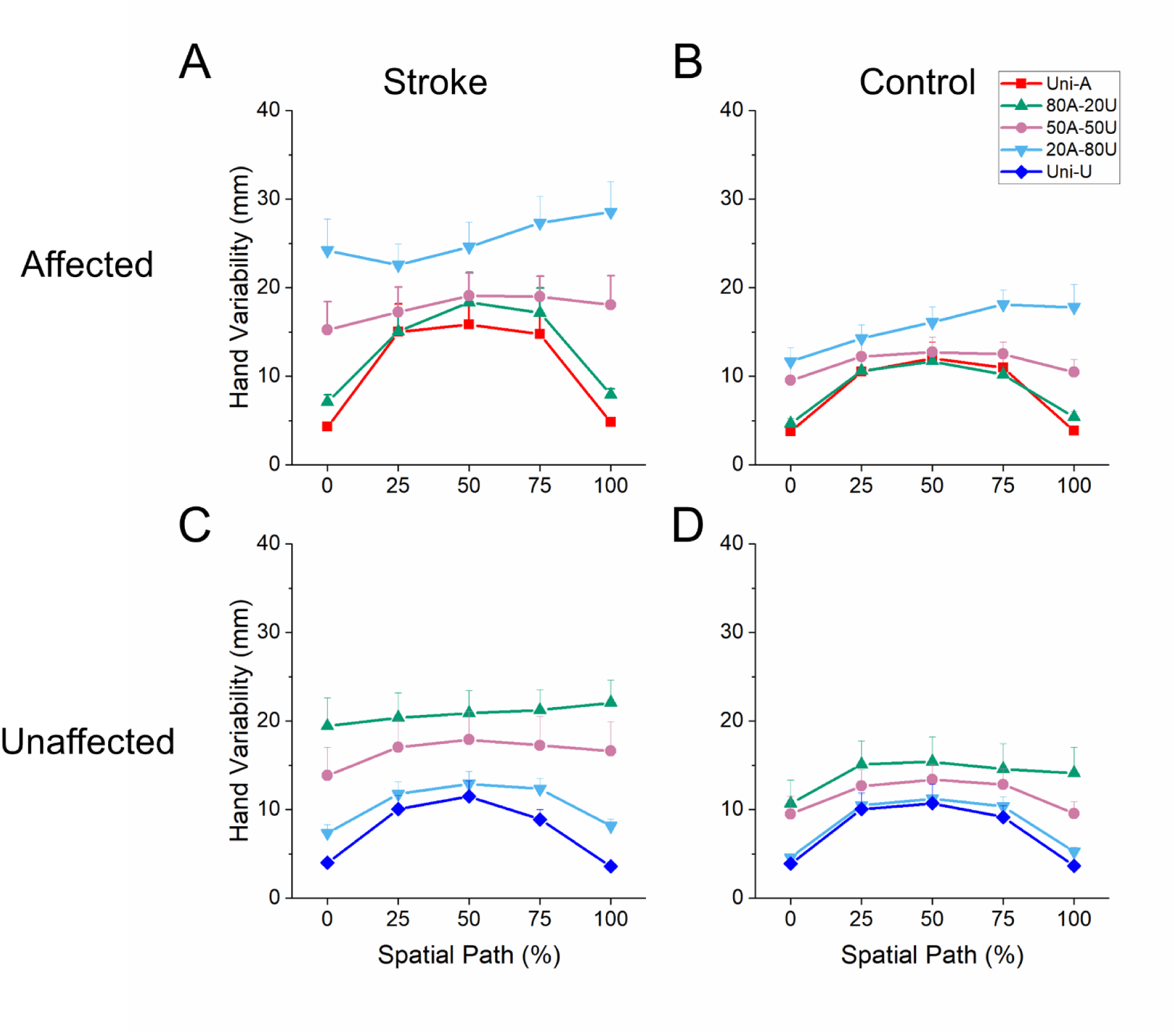
Variability of the affected hand in (A) stroke and (B) control groups. Only four of the five conditions are shown because the affected arm was not used in the Uni-U condition. Participants in both groups showed higher variability as the contribution of the arm to cursor motion decreased. Variability of the unaffected hand in (C) stroke and (D) control groups. Again, only four of the five conditions are shown because the unaffected arm was not used in the Uni-A condition. Similar to the affected arm, participants in both groups showed higher variability in the unaffected arm as the contribution of the arm to cursor motion decreased.

### 5.3 Correlation between left and right arms

Similar to the cursor variability, the analysis of the correlation between the two arms during the movement was modulated along the path and was influenced by condition. There was a main effect of phase, F(2.22,44.40) = 96.018, p<.001, and a main effect of condition, F(2, 40) = 6.141, p = .005. Post-hoc analysis of the phase effect showed that similar to the variability, the correlation was more negative during the start and end of the movement, but was weaker in the middle of the movement. Post-hoc analysis of the condition effect showed that the 50A-50U condition showed a more negative correlation than the 80A-20U condition.

**Figure 5.**
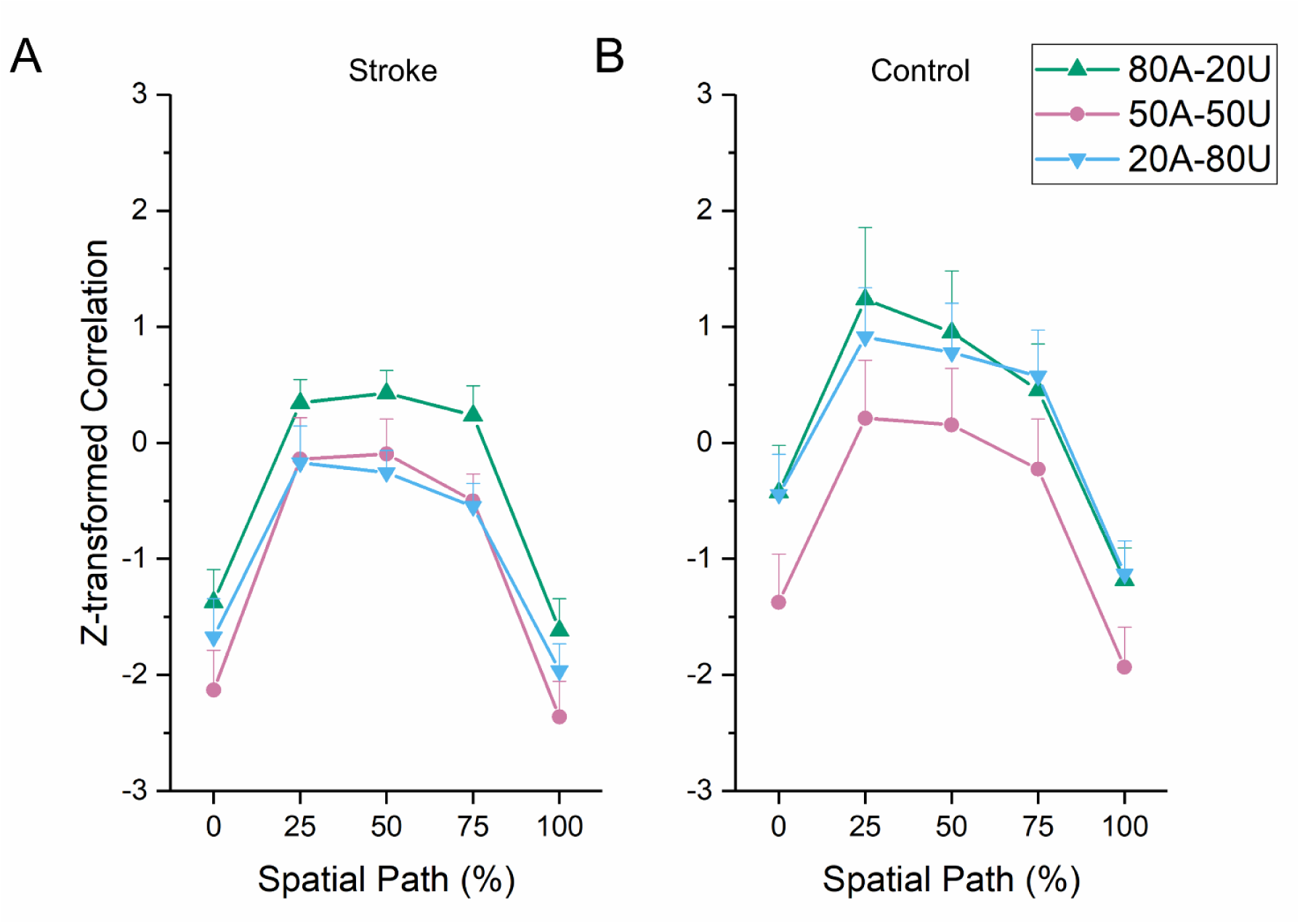
Z-transformed correlations between the affected and the unaffected arms at different points along the spatial path in (A) stroke and (B) control groups. Only the 3 bimanual conditions are shown (as the 2 unimanual conditions did not involve motion of both arms). Again, both the stroke and control groups showed similar modulation of the correlation along the path in all conditions, with a strong negative correlation at the start and end of the trajectory and a weaker correlation near 50% - i.e. roughly halfway along the distance between the home position and the target.

## 6 DISCUSSION

The purpose of the current study was to examine how stroke survivors can modulate the variability in the affected and unaffected arms. Using a bimanual shared task where we could alter the task demands by altering the weighting (i.e. the degree to which each arm contributed to the task), we examined if the variability in each arm was sensitive to the change in these task demands. Overall our results showed that similar to the control group, stroke survivors were able to modulate the variability in each arm depending on the task demands. Movement times in stroke survivors were longer than controls – yet, in terms of the variability the stroke survivors generally showed higher variability, indicating that the results are not the consequence of a speed-accuracy tradeoff.

When we considered the spatial variability of the cursor (i.e., the object that was controlled in both unimanual and bimanual tasks) in the direction perpendicular to the reach, we found that consistent with prior studies (Khan et al., 2002), the variability showed a distinct pattern during the reach; this was characterized by low variability at the start and end of the movement, and peak variability occurring at about 50% into the movement. This variability profile of the cursor was also reflected in the variability of the arm in unimanual tasks because there was no redundancy in the unimanual task. However, when we considered the bimanual tasks (which had redundancy), we found that variability profile of the cursor did not always reflect the variability of the arms. Instead, we found that the arm variability was modulated in bimanual tasks such that if the weighting of the arm to cursor motion was low, the variability of the arm increased, and did not show the same profile as the cursor variability. On the other arm, when the weighting of the arm to cursor motion was high, the variability of the arm typically decreased and started resembling the same profile as the cursor variability. While the affected arm in stroke survivors had higher variability overall, this same pattern of results was seen in both the unaffected and the affected arms, and in age-matched controls as well.

This raises the question – if the arm variability increased in a particular condition, why did the cursor variability not increase? The answer to this question is addressed by our correlation analyses that examined if the variability in the two arms was related. Our correlation results showed a strong negative correlation between the two arms, indicating a strong compensatory action (i.e. when one hand moved, the other hand moved in the opposite direction). As has been shown in prior studies (Latash et al., 2002; Todorov and Jordan, 2002; Diedrichsen, 2007), the presence of a strong negative correlation between the two arms allows the individual arms to be more variable, while still controlling the cursor variability. Interestingly, these results were not different in the stroke survivors and the control group suggesting that in spite of the higher variability in the affected arm, this coordination between the two arms was not affected by the stroke, at least in stroke survivors with mild to moderate hemiparesis. These results are consistent with previous results showing that the ability to coordinate multiple joints of the same arm to stabilize reaching performance are preserved in individuals with mild or moderate hemiparesis (Reisman and Scholz, 2003).

From a theoretical standpoint, the fact that variability can be modulated by the task is consistent with the idea that the measured variability in stroke survivors is not entirely ‘noise’, but can be actively controlled (Newell and Slifkin, 1998; Harbourne and Stergiou, 2009). In particular, the results are consistent with an optimal feedback control view of motor redundancy (Todorov and Jordan, 2002) for two reasons. First, the fact that variability increases in the arm that contributes less to cursor motion is consistent with the idea that the motor system does not ‘waste’ effort in reducing variability of effectors that do not contribute significantly to the task. Moreover, the fact that the correlation starts off negative at the start of the movement, becomes weaker during the middle portion of the reach, and becomes negative again during the end of the reach, suggests that these correlations are likely mediated through visual feedback.

Finally, our results also have implications from a rehabilitation standpoint. Previous studies on bimanual therapy have focused on temporal and spatial movement parameters (McCombe Waller et al., 2008), or on the phase relation between the arms when the two arms move simultaneously (Sleimen-Malkoun et al., 2011). Here, we show that in shared bimanual tasks, there is modulation of variability in each arm, and a coupling that emerges between the two arms. The implications of this coupling for therapy are unclear at the moment – one interpretation is that the bimanual shared task here facilitated a ‘cooperative’ strategy between the two arms, which potentially could be an important target for rehabilitation (Sainburg et al., 2013). However, on the other hand, we also found that the affected arm became more variable (i.e. control was “sloppier”) when the unaffected arm was involved. Therefore an alternative interpretation is that the bimanual shared tasks encouraged ‘compensatory movements’ from the unaffected arm to stabilize the task. Whether these compensatory behaviors are adaptive or maladaptive in terms of ultimately improving motor function of the affected arm in the long-term still remains an open question.

## 7 ACKNOWLEDGMENT

The present study was funded in part by the MSU-Sparrow Center for Innovation and Research. We also thank Julie Pieciak and Ten-Niah Kinney for assisting with data collection.

